# Quantitative comparison of methodologies for translation site imaging in living cells

**DOI:** 10.1101/2025.10.29.685314

**Authors:** Agata D. Misiaszek, Esther Griesbach, Egle Jaugaite, Aurelio Ortale, Jan Eglinger, Tobias Hochstoeger, Jeffrey A. Chao

**Affiliations:** Friedrich Miescher Institute for Biomedical Research, 4056 Basel, Switzerland

## Abstract

Single-molecule imaging of translation sites in living cells has enabled the dynamics of protein synthesis to be investigated with high spatial and temporal resolution. These methodologies utilize the interaction between a multimerized epitope tag and its cognate fluorescent nanobody to detect nascent polypeptides as they emerge from the ribosome. Here, we present a systematic comparison of current methodologies and determine that the ALFA-tag reduces perturbations of mRNA expression and increases the fluorescent signal of translation sites.

## Main

Fluorescence microscopy methodologies have enabled visualization of translation in living cells^1,2^. Among these approaches, the SunTag system^3^ combines a tandem array of optimized GCN4 epitopes with a GFP-tagged single-chain variable fragment antibody (scFv-GFP)^4^ (Supplementary Tab.1) to amplify nascent peptide signals, allowing quantification of translation dynamics with single-molecule resolution^5–8^. SunTag-based imaging has provided insights into translational regulation across diverse contexts, including subcellular localization^9–11^, stress responses^12–14^, RNA stability ^15^, and frameshifting^16,17^. While the SunTag methodology has been the most widely adopted technique, alternative strategies have used orthogonal multimerized epitope tags based on HA epitopes and the ALFA-tag (Supplementary Tab.1)^18–20^. Additionally, the HA epitopes were grafted into the loops of a nonfluorescent GFP scaffold known as a spaghetti monster, which provides a structured scaffold compared to linear arrays of epitopes^21^.

To systematically compare these methodologies, we used a doxycycline-inducible HeLa cell line with a defined genomic locus for site-specific integration of reporter genes to reduce cell-to-cell expression heterogeneity resulting from transient transfection^13,22–25^. Reporter mRNAs were generated with coding sequences that contained a linear array of 18x GCN4 epitopes (SunTag), a spaghetti monster with 18xGCN4 epitopes (smGCN4), a spaghetti monster with 18xHA epitopes (smHA), a spaghetti monster with 18xALFA-tag (smALFA) or EGFP, which was chosen because it is similar in size to a spaghetti monster. All reporter constructs also contained 24xMS2 stem-loops in the 3′ untranslated region (UTR) to enable detection of single mRNAs (Fig 1A). The HeLa cells stably expressed the cognate fluorescent nanobodies and MS2 coat proteins (MCP) and cells with appropriate expression levels for single-molecule imaging were isolated by fluorescence activated cell sorting.

**Figure 1.**
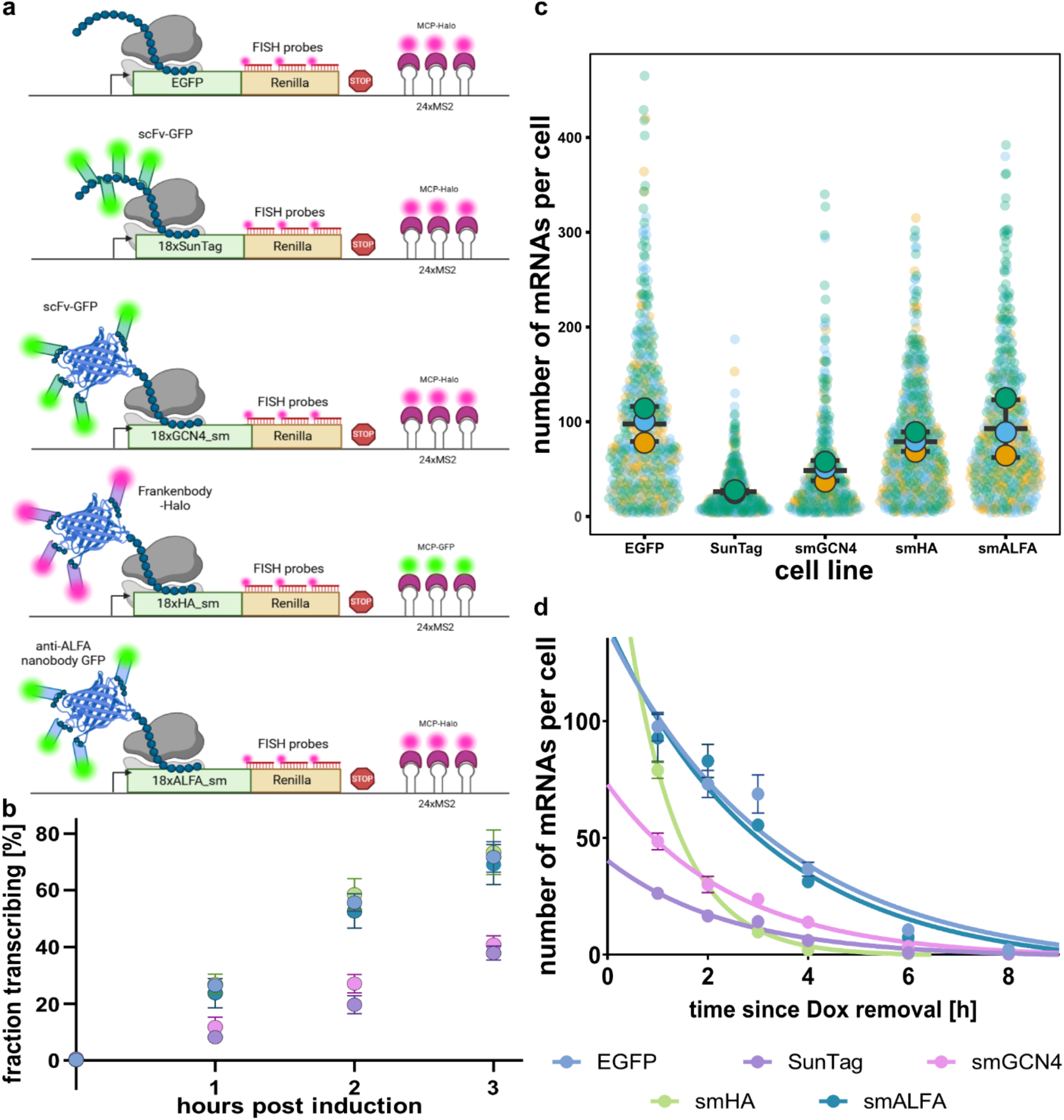
Comparison of RNA expression and stability in mRNA reporter constructs. **a**, Schematic of reporter constructs and components used for live-cell single-molecule imaging. sm – spaghetti monster; scFv – single-chain variable fragment antibody; FISH – fluorescence in situ hybridization; MCP – MS2 coat protein. **b**, Fraction of cells actively transcribing after doxycycline induction. Error bars represent standard deviation (SD); mean of three independent experiments performed on separate days. **c**, Number of mRNA spots per cell after 1.5 h induction and removal of doxycycline for 1h. Independent replicates were collected on separate days. Cells analyzed: EGFP *n* = 438, SunTag *n* = 389, smGCN4 *n* = 289, smHA *n* = 433, smALFA *n* = 362. Data shown as mean ± SD. **d**, mRNA decay of reporter constructs. Cells were induced with doxycycline for 1.5 h, washed for 30 min, and imaged at indicated times post-washout. Trendlines were fitted using a non-linear one-phase decay model. Error bars indicate standard error of the mean (SEM); mean of three independent experiments shown.

To determine if the epitope tagging altered transcription of the reporter mRNAs, we quantified the fraction of cells that contained mRNAs after doxycycline induction (Fig. 1b, Supplementary Fig. 1) The fraction of induced cells increased overtime for all five reporter mRNAs, however, at all timepoints the number of cells containing SunTag or smGCN4 transcripts was markedly lower than the EGFP, smHA, and smALFA reporters. At three hours after doxycycline induction, ∼70% of cells contained EGFP, smHA or smALFA transcripts, while only ∼40% of cells expressed SunTag or smGCN4 transcripts. We also quantified the number of transcripts per cell and found that EGFP (97.6±6.1) and smALFA (92.8±10.1) reporter mRNAs had similar mRNA levels while smHA (79.0±3.4) had slightly lower numbers and SunTag (26.2±0.5) and smGCN4 (48.5±3.5) had substantially less transcripts. smGCN4 transcripts were expressed at higher levels than SunTag suggesting that the spaghetti monster scaffold may help to mitigate some of the negative effects of the GCN4 epitopes placed in a repetitive linear array. To determine, if the epitope tags also affected mRNA stability, we washed out doxycycline to stop transcription and quantified the number of transcripts per cell at successive time points (Fig. 1e; Supplementary Figs. 2-3). The EGFP and smALFA reporter mRNAs exhibited similar apparent half-lives (t½ = 2.58 h and 2.36 h, respectively) and the SunTag and smGCN4 reporters decayed faster (t½ = 1.8 h). Interestingly, smHA mRNAs were the least stable (t½ = 1 h) despite being transcribed at a higher level than SunTag and smGCN4 transcripts. In these experiments, the smALFA was found to have minimal effects on transcription and mRNA stability, while the GCN4 epitopes in either the linear or spaghetti monster array reduced transcription and the smHA affected mRNA stability.

Next, we compared the performance of the epitope tags for live-cell translation site imaging. Initially, we fused GFP to all of the nanobodies, however, we observed that the HA-Frankenbody-GFP performed poorly for imaging translation sites, so we generated an MCP-GFP with HA-Frankenbody-Halo cell line. All reporters mRNAs were induced to achieve comparable transcript numbers (SunTag and smGCN4, 1.5 h; smHA and smALFA, 1 h) (Fig. 2a) to ensure that nanobody levels were not depleted at the time of imaging. We found that ∼80% of SunTag, smGCN4 and smALFA reporter RNAs were actively being translated, consistent with previous observations^12^, whereas ∼57% of smHA RNAs were undergoing translation and with higher variability between biological replicates (Fig. 2b; Supplementary Fig. 4a). Translation site intensity measurements revealed that smALFA foci were ∼1.6-fold brighter than SunTag and smGCN4 translation sites and ∼2.7-fold brighter than smHA foci (Fig. 2c; Supplementary Fig. 4b). While linear GNC4 epitopes in the SunTag array have previously been found to be saturated with scFv-GFP^3,8^, our results suggests that the smALFA achieves higher nanobody occupancy when the nanobodies are expressed at similar levels (Supplementary Fig. 4c,d).

**Figure 2.**
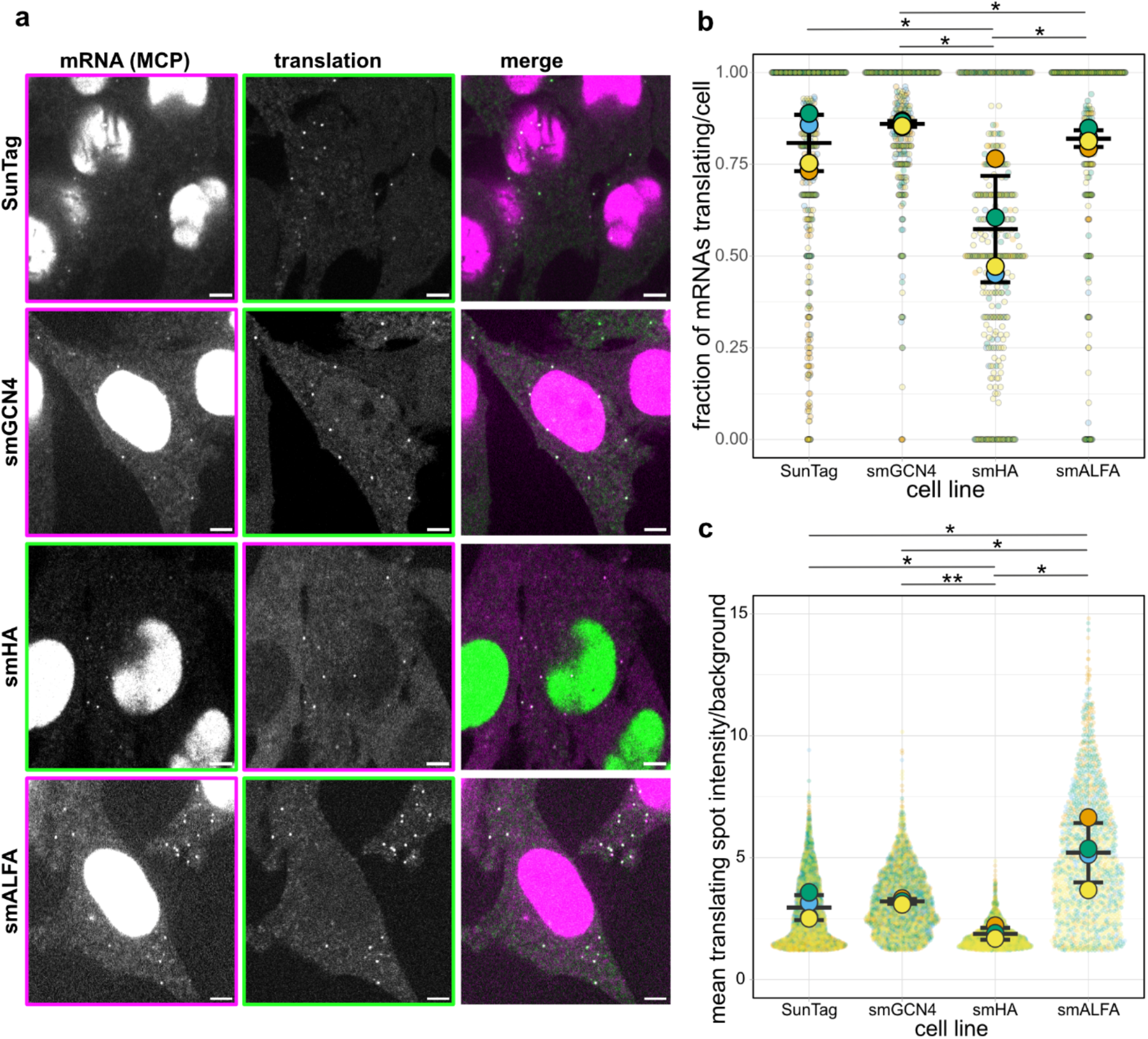
Live-cell imaging of reporter cell lines. **a**, Representative images of each reporter cell line. Scale bar: 10 μm. **b**, Fraction of mRNA molecules undergoing translation in each cell line. **c**, Translation site intensity normalized to background. In **b** and **c**, data are shown as mean ± SD from four independent replicates collected on separate days. Statistical significance: *p* < 0.05 (*), *p* < 0.001 (**). Number of cells analyzed: SunTag *n* = 524, smGCN4 *n* = 369, smHA *n* = 441, smALFA *n* = 343.

In conclusion, we have performed a systematic comparison of the current methods for single-molecule translation-site imaging. We find that the smALFA has minimal effects on transcription and mRNA stability, while SunTag and smGCN4 impact transcription and smHA reduces mRNA stability. Additionally, though all of the approaches enabled imaging of translation sites in living cells, we found that the smALFA generated the brightest translation sites. It should be noted, however, that while the HeLa cell line used in this study enabled a quantitative comparison between the methodologies, the effects of the epitope tags could be different in other cell lines or model systems. It is critical that effect of the epitope tags be considered during experimental design to minimize their impact on the underlying biological process being studied.

## Methods

### Cell culture

This study used HeLa-11ht cells previously described^25^ containing an Flp-RMCE (recombinase-mediated cassette exchange) site for single-copy integration at a defined genomic locus and the reverse tetracycline-controlled transactivator (rtTA2S-M2) for doxycycline-inducible expression of the inserted gene. Cells were cultured in Dulbecco’s Modified Eagle Medium (DMEM) with 4.5 g/L glucose, 100 U/mL Penicillin, 100 μg/mL Streptomycin, 4 mM L-Glutamine, 10% fetal bovine serum (FBS) at 37ºC and 5% CO2.

### DNA constructs

All reporter plasmids were cloned by Gibson assembly^26,27^ using the SunTag-Renilla-MS2 plasmid as a backbone^13^ (Addgene #119945) and replacing the existing SunTag cassette with different inserts. For EGFP-Renilla-24xMS2 plasmid, EGFP was cloned from a G3BP1-GFP plasmid (Addgene #119950); for the sm-18xGCN4-Renilla-24xMS2 plasmid, sm-18xGCN4 was cloned from a synthesized fragment (Genwiz); for the sm-18xHA-Renilla-24xMS2 plasmids and sm-18xALFA-Renilla-24xMS2 plasmids the sm-18xHA and sm18xALFA inserts were cloned from synthesized fragments (Genscript).

### Cell line generation

NLS-stdMCP-stdHalo / NLS-stdMCP-stdGFP / HA-Frankenbody-Halo / ALFA-Nanobody-GFP were stably integrated into cell lines using a lentiviral system^28^ in biosafety level 2 conditions. Briefly, lentivirus was produced by transfecting HEK293T cells with pHAGE lentiviral backbone and accessory plasmids VSV-G, Gag/Pol, Tat, Rev using Fugene 6 (Promega, E2691). The supernatant was harvested for the next three days before being centrifuged, filtered through a 0.45 μM filter (Milipore, SLHV033RS), and concentrated with LentiX concentrator (Clontech, 631232). HeLa cells were infected for 6 days, passaged, and Fluorescence-activated Cell Sorting (FACS) was performed to screen for cells with appropriate expression levels of NLS-stdMCP-Halo for single-molecule imaging.

EGFP-Renilla-MS2 plasmids were integrated into HeLa-11ht cell line stably expressing NLS-stdMCP-stdHalo. smGCN4-Renilla-MS2 and SunTag-Renilla-MS2 plasmids were integrated into a HeLa-11ht cell line stably expressing GFP-tagged single-chain antibodies (scFv-GFP) against GCN4^23^ (Addgene #104998). NLS-stdMCP-stdHalo was then added using lentiviral system as described. smHA-Renilla-MS2 plasmid was integrated into HeLa-11ht cell line expressing HA-Frankenbody-Halo-GB1 and NLS-stdMCP-stdGFP. smALFA-Renilla-MS2 plasmid was integrated into HeLa-11ht cell line expressing ALFA-Nanobody-sfGFP and NLS-stdMCP-stdHalo.

All reporters were integrated via RMCE as described previously^25^. Specifically, on the first day, 3 ×10^5^ HeLa cells were seeded into a 6-well plate. On day 2, cells were transfected with the reporter mRNA plasmid (2 μg) and pCAGGS-FLPe-IRESpuro plasmid (2 μg, Addgene #20733) using Lipofectamine 2000 (Invitrogen) according to the manufacturer’s protocol. The subsequent day, 2 μg/mL puromycin (Invivogen, ant-pr-1) was added to the culture medium to select for transfected cells. On day 5, puromycin was removed and 50 μM ganciclovir (Sigma-Aldrich, G2536) was added to the media for the next 10 days to select cells that had undergone successful RMCE. Cell lines with appropriate levels of fluorescent proteins were then isolated by FACS and the reporter integration was confirmed with microscopy.

### RNA induction measurement

Cells were seeded by transferring 100µl of cell suspension at the density of 7.0 × 10^4^ cells/well into an empty, poly-lysine-coated 96-well glass-bottom dish (Cellvis P96-1.5N) using an Integra Assist Plus equipped with an Integra VOYAGER - 300µl – 8 channel pipette and cultured for 48h. Reporter mRNA transcription was induced by addition of 1 μg/mL doxycycline for indicated by the time-point length (0h – no doxycycline added, 1h, 2h and 3h). Cells were fixed by direct addition of 2x volume of 8% paraformaldehyde (Electron Microscopy Sciences) in 2x PBS directly to cell culture medium (final concentration 4% PFA and 1x PBS) for 10 min. Cells were permeabilized in 70% ethanol overnight at 4°C.

### RNA decay measurement

Cells were seeded on high-precision glass coverslips (170 µm thickness, 18 mm diameter; Paul Marienfeld GmbH) in 12-well plates at a density of 8.0 × 10^4^ cells/well and cultured for 48 h.

Reporter mRNA transcription was induced by treating cells with doxycycline for 1.5 h. To stop transcription, doxycycline was removed by washing the cells three times with culture medium. At designated time points starting 30 min after doxycycline removal, cells were washed twice with PBS, fixed in 4% paraformaldehyde (Electron Microscopy Sciences) in PBS for 10 min, and permeabilized in 70% ethanol overnight at 4°C.

### Fluorescence in-situ hybridization

smFISH was performed as described previously^24^. Probes targeting the Renilla luciferase open reading frame (Supplementary Table 2) were enzymatically labeled with Atto-633 using terminal deoxynucleotidyl transferase^29^. Briefly, cells were washed twice with PBS, pre-blocked in warm wash buffer (2× SSC, 10% formamide) for 5–10 min, and hybridized in hybridization buffer (125 nM smFISH probes targeting the Renilla coding sequence, 2× SSC, 10% formamide, 10% dextran sulfate (Millipore)) for 4 h at 37 °C in the dark. Cells were then washed in pre-warmed wash buffer for 30 minutes at room temperature. Finally, coverslips were washed twice with PBS and mounted in ProLong Gold antifade reagent with 4,6-Diamidino-2-phenylindole dihydrochloride (DAPI) (Invitrogen) for the RNA decay experiment. For the RNA induction experiment, 2 μg/mL DAPI (Sigma-Aldrich, D9542) was added for the last PBS wash and no mounting medium was used.

For the RNA induction experiment, high-throughput imaging was performed on a Yokogawa (CellVoyager 7000S), an automated spinning disk microscope with enhanced CSU-W1 spinning disk (Microlens-enhanced dual Nipkow disk confocal scanner), an Olympus 60xW UPLS APO objective (NA = 1.2), and a Neo sCMOS camera (Andor, 2,560 × 2,160 pixels). Obtained images were recorded with 0.108 µm/pixel.

For the RNA decay experiment, images were acquired using a Zeiss AxioObserver7 inverted microscope with a Yokogawa CSU W1-T2 spinning disk confocal unit, Plan-APOCHROMAT 100x/ 1.4 NA oil objective, sCMOS camera, and X-Cite 120 EXFO metal halide light source. 21 z-stacks were collected at 0.21 µm steps. Exposure times were 600 ms for Atto-565 and 100 ms for DAPI. Images were acquired with the pixel size of 0.107 μm. Cells were selected for imaging randomly using the DAPI channel.

### smFISH image analysis for RNA induction experiment

For analysis of the data from the RNA induction experiment, cells were segmented using a nuclei-anchored 3D workflow. Nuclei (stained with DAPI) were segmented in 3D using custom script available on GitHub. The cytoplasm was segmented on a 2D maximum intensity projection (MIP) of DAPI after median filtering using a watershed mask built from opening, area opening, and hole filling. Candidate mRNA spots were detected in 3D by a scale-matched Laplacian-of-Gaussian (σ ≈ λ/(2·NA·√2) converted to pixels) with h-maxima (contrast h), then refined by subpixel 3D Gaussian fits (background + anisotropic Gaussian yielding x,y,z and σx,σy,σz). Spots were classified as nuclear/cytoplasmic by sampling the 3D nucleus label and 2D cell mask at fitted centroids and filtered for symmetry. Cells with ≤3 spots were classified as uninduced based on spot detection in cells in the absence of doxycycline.

### smFISH image analysis for RNA decay

For each field of view, two z-stacks were acquired (smFISH and DAPI nuclear stain) and converted to 2D MIPs. Pixel size and units were read from TIFF UIC/OME tags and used downstream for scale-aware filtering. Cytoplasm was segmented on the smFISH MIP with Cellpose^30^ using the cyto3 model (Gaussian pre-smoothing σ = 1; nominal object diameter parameter set to 250 px), and nuclei were segmented on the nuclear MIP with Cellpose (automatic size estimation). We retained only nuclei whose bounding boxes were fully contained in the image to avoid edge artifacts; “valid cells” were defined as cytoplasmic masks overlapping these accepted nuclei. smFISH spots were detected with the big-fish pipeline^31^. A Laplacian-of-Gaussian filter was applied with σ estimated from the physical pixel size and an expected spot radius of 150 nm; local maxima were found, and an automated intensity threshold was used to call spots. For each detection we drew a labeled disk of radius = 2 px to create a dot mask and extracted standard properties (centroid, area, min/max/mean and total intensity). Background was estimated from the cytoplasmic compartment after explicitly removing spot pixels; the signal-to-background ratio (SBR) for each spot was defined as mean spot intensity divided by this background mean. Unless noted otherwise, only spots with SBR ≥ 3 were retained. Spots were assigned to nucleus or cytoplasm by sampling the corresponding masks at spot centroids. Manual quality control was performed afterwards. For every valid cell we reported compartment-resolved counts (“spots in nucleus”, “spots in cytoplasm”); cells with ≤3spots were considered uninduced and kept to preserve denominators in summaries. Aggregation was performed across samples within each timepoint.

### Live-cell imaging

For live-cell imaging, NLS-stdMCP-stdHalo or NLS-stdMCP-stdGFP was stably integrated into cells expressing scFv-GFP / HA-Frankenbody-Halo / ALFA-Nanobody-sfGFP and SunTag, smGCN4, smHA or smALFA reporter constructs, as described above. 3.5×10^4^ cells were seeded in 35 mm Ibidi glass-bottom dishes (ibidi, 81158) and grown for 48 h. NLS-stdMCP-stdHalo / HA-Frankenbody-Halo was labeled with JF646 HaloTag ligand (100nM) for 20 minutes (HHMI Janelia Research Campus^32^). Cells were washed 3 times in DMEM to wash out unbound ligand. Reporter mRNA expression was induced with 1 μg/mL doxycycline for 1 hour for smALFA and smHA reporters and 1.5 hours for SunTag and smGCN4 reporters to ensure comparable spot counts. Cells were kept at 37°C and 5% CO_2_ during the entire imaging session.

Imaging was performed as described previously^12,14,24^ using an inverted motorized stand Nikon Ti2-E Eclipse microscope equipped with a Yokogawa CSU W1 with Dual camera T2 spinning disk confocal scanning unit, two iXon-Ultra-888 (Andor) cameras and a CFI P-Apo Lambda 100x 1.45 NA oil objective. 488 iBeam Smart (Toptica Photonics) and 639 iBeam Smart (Toptica Photonics) lasers were employed for illumination, together with a VS-Homogenizer (Visitron Systems GmbH). Images were acquired using VisiView software (Version 5.0.0.14) in a single plane at frame rates of 20 Hz (50 ms exposure). Images were acquired with the pixel size of 0.134 μm. Images of 0.5 μm TetraSpeck beads (Thermo Fisher Scientific, T14792) were acquired to correct for camera misalignment.

### Live-cell image analysis

Image analysis was performed as described^12^. Briefly, frames 5 to 14 of each movie (500 ms) were selected and any offset between the two cameras was corrected using TetraSpeck fluorescent beads acquired on each imaging day and FIJI descriptor-based registration plugin in affine transformation mode. Subsequently, if necessary, fine correction for each dish individually was performed using the FIJI translate function run in a custom macro.

Using the KNIME analytics platform and a custom-build data processing workflow, regions of interest (ROIs) were manually annotated in the mRNA channel excluding any bias in selection attributable to the translational state of the cell. Then, spots corresponding to single mRNAs were tracked using TrackMate integrated in KNIME, using the “Laplacian of Gaussian” detector with an estimated spot radius of 200-nm and subpixel localization. Detection thresholds were adjusted by taking the mean signal from the ROIs multiplied by 1.25. For particle linking, the parameters linking max distance (600 nm), gap closing max distance (1200 nm), and gap closing max frame gap (2) were used. To assay whether an mRNA is translating, the mean intensity of the translation channel was measured at the coordinates of each mRNA spot and quantified as FC/ROI background intensity. A cutoff of <1.5 fold/background was determined to classify an mRNA as non-translating.

### Western Blotting

Western blotting was performed as described^12^. In summary, for protein extraction, cells were harvested by trypsinization and lysed in radioimmunoprecipitation assay buffer (150 mM NaCl, 50 mM tris, 0.1% SDS, 0.5% sodium deoxycholate, and 1% Triton X-100) supplemented with 1× protease inhibitor cocktail (Thermo Fisher Scientific, 78440) and SuperNuclease (Sino Biological). Cell lysate was centrifuged at 12,000 rpm for 10 min to remove cell debris, and the supernatant was loaded on a 4 to 15% gradient gel using loading buffer supplemented with 100 mM dithiothreitol. Following SDS–polyacrylamide gel electrophoresis (SDS-PAGE), proteins were transferred onto nitrocellulose membranes by semi-dry transfer (Trans-Blot Turbo) and blocked in 5% bovine serum albumin-TBST buffer (tris-buffered saline supplemented with 0.1% Tween 20) for 1 hour at room temperature (RT). Primary antibodies (mouse anti-GFP (Abcam, ab183734) and as used as a loading control goat anti-GAPDH (Invitrogen, Pa1-9046)) were incubated overnight at 4°C in Intercept blocking buffer (LI-COR) supplemented with 0.1% Tween 20. The next day, the membrane was washed three times in TBST and incubated for 1 hour at RT with the fluorescent secondary antibodies (IRDye 680RD Goat anti-Mouse IgG (LI-COR, 926-68070) and IRDye 800CW Goat anti-Rabbit IgG (LI-COR, 926-32211)) diluted 1:10,000 in Intercept blocking buffer with 0.1% Tween 20. Following three washes in TBST, membranes were transferred to phosphate-buffered saline (PBS), and antibody fluorescence was detected at 700 and 800 nm using an Odyssey infrared imaging system (LI-COR).

### Statistical analysis

For smFISH decay experiment number of inducing cells was normalized to the 1h timepoint. All statistical analysis was performed on the means of the independent replicates. Statistical analysis for the smFISH experiments was performed using GraphPad Prism 10.4.1. For the RNA decay data, one phase decay non-linear fit curves were automatically fitted. For live-cell imaging, data was plotted and the statistical significance was calculated using SuperPlotsOfData^33^.

## Metadata

### Code availability

All analysis workflows generated in this study have been made publicly available on GITHUB.

### Data availability

All data will be made available on request.

## Acknowledgments

We would like to thank the Facility for Advanced Imaging and Microscopy at the FMI, especially S. Reither, for their support with microscopy and image analysis and H. Kohler from the Cell Sorting facility for support in cell line selection. We are grateful to B. T. Eichenberger for his help with cell line generation and analysis of the initial data, V. Bhaskar assisted with cloning of the smGCN4 plasmid and F. Naef and C. Gobert for helpful discussions concerning smHA. This work was supported by the Novartis Research Foundation (J.A.C), a Swiss National Science Foundation Sinergia grant (CRSII5-205884) (J.A.C), the SNF-NCCR RNA & Disease network (51NF40-205601) (J.A.C) and a EMBO postdoctoral fellowship (ALTF_179-2023)(A.D.M).

## Author contributions

ADM, TH and EG created the cell lines. ADM, TH and EJ carried out the experiments. ADM, AO and JE performed the data analysis. The manuscript was written with input from all authors.

---

**Supplementary Table 1.** Main characteristics of the compared translation site imaging systems.

| Cell line | Insert in the open frame | Sequence of the repetitive epitope | Tag size [kDa] | Nanobody | Nanobody size [kDa] | Nanobody affinity |
| --- | --- | --- | --- | --- | --- | --- |
| EGFP | EGFP | - | 26.8 | - | - | - |
| SunTag | 18xGCN4 (linear) | EELLSK<br>NYHLEN<br>EVARLK<br>K | 65.3 | SunTag single-chain variable fragment (scFv) | 25.76 | 44 pM |
| smGCN4 | 18xGCN4 (spaghetti monster) | EELLSK<br>NYHLEN<br>EVARLK<br>K | 75.1 | SunTag single-chain variable fragment (scFv) | 25.76 | 44 pM |
| smHA | 18xHA (spaghetti monster) | YPYDVP<br>DYA | 51.8 | HA-Frankenbody | 27.6 | 14.7 nM |
| smALFA | 18xALFA (spaghetti monster) | PSRLEE<br>ELRRRL<br>TEP | 68.8 | ALFA-Nanobody | 13.45 | 26 pM |

**Supplementary Figure 1.**
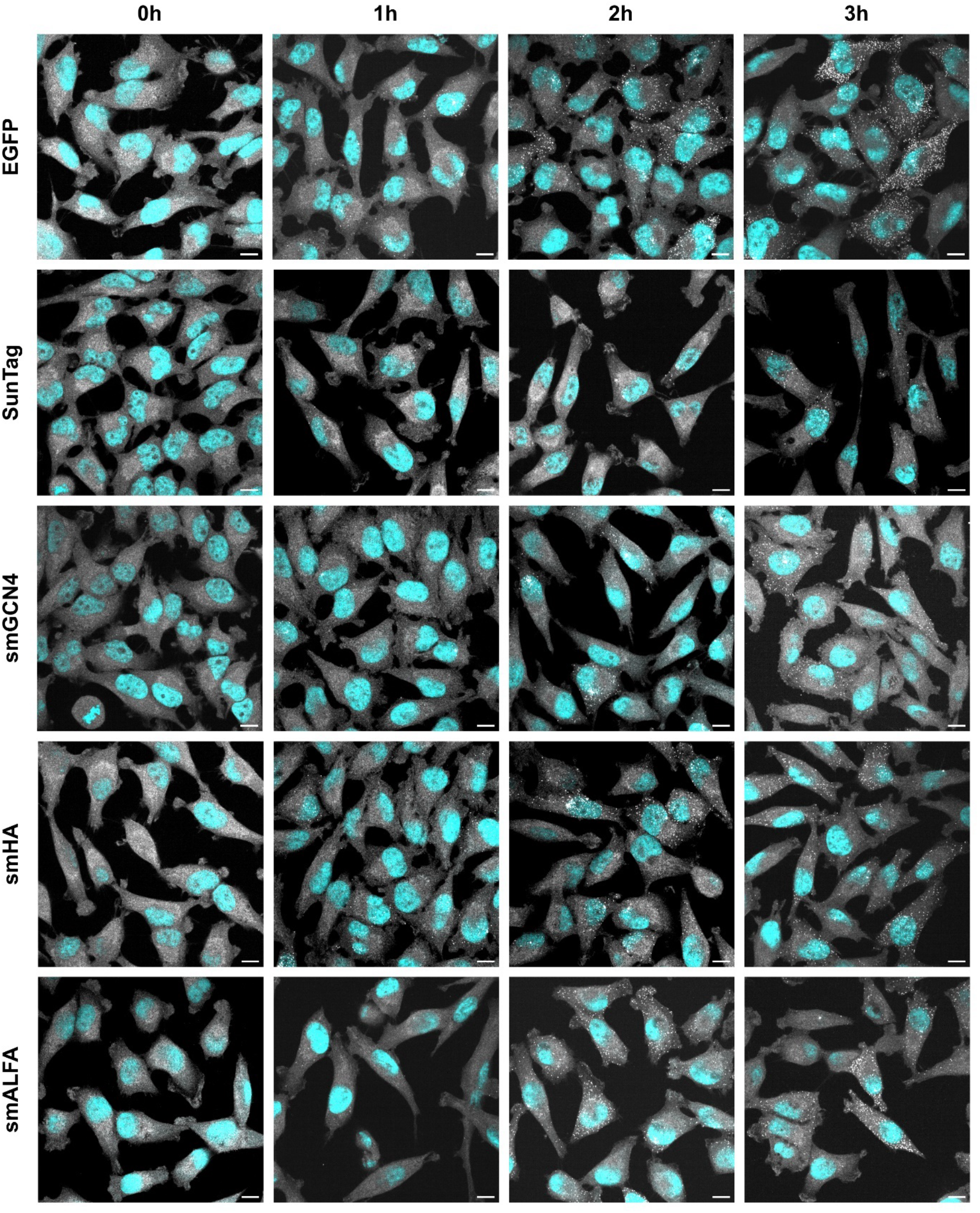
Representative images used for quantification of the fraction of cells inducing mRNA transcription in respective cell lines. A single plane from the center of the cell is shown. Cyan – DAPI, grey – mRNA. Scale bar: 10 μm

**Supplementary Figure 2.**
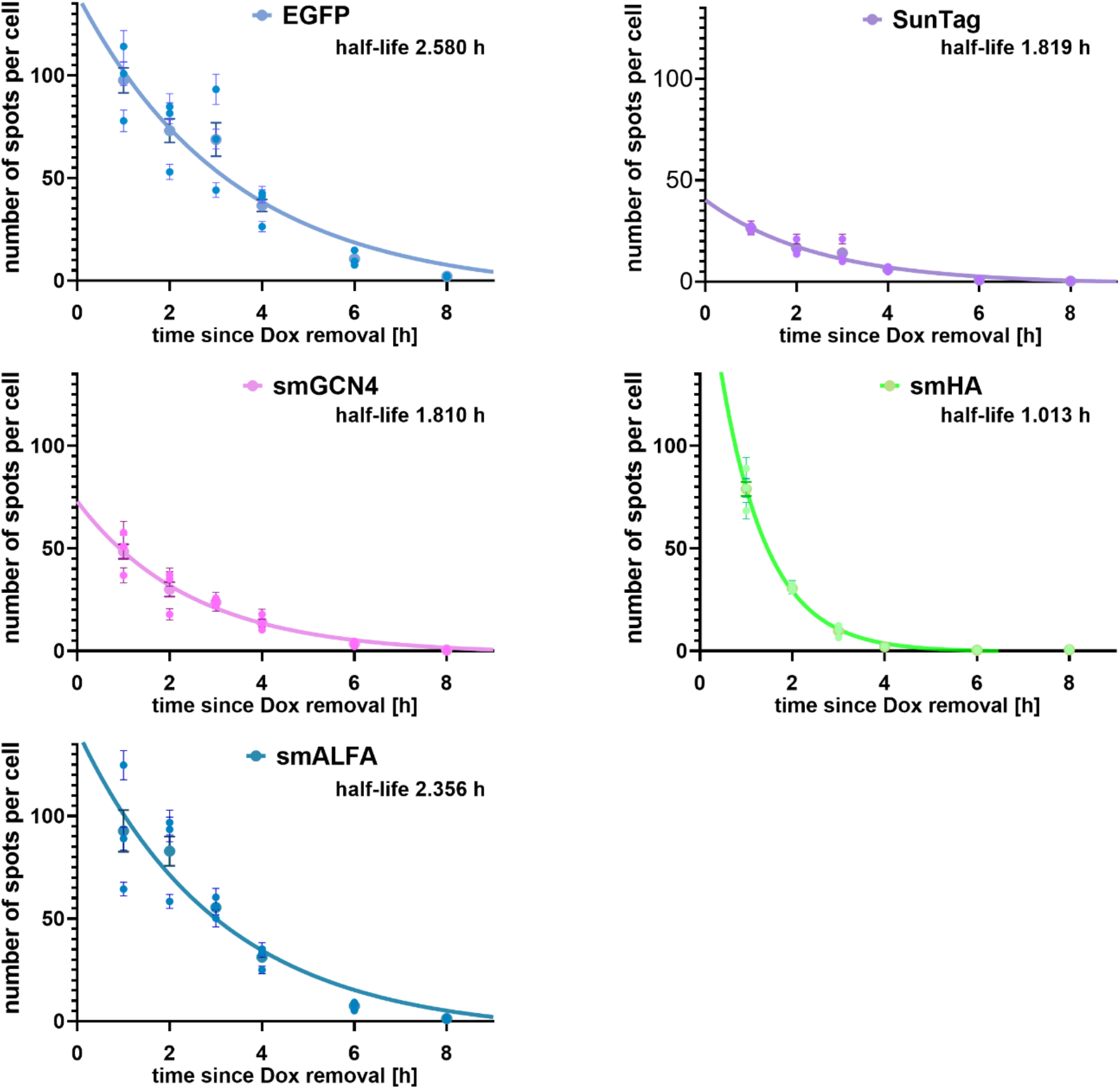
mRNA decay curves for each reporter cell line. Three independent replicates are shown separately, with error bars representing standard deviation. Half-lives were calculated from non-linear one-phase decay curve fits.

**Supplementary Figure 3.**
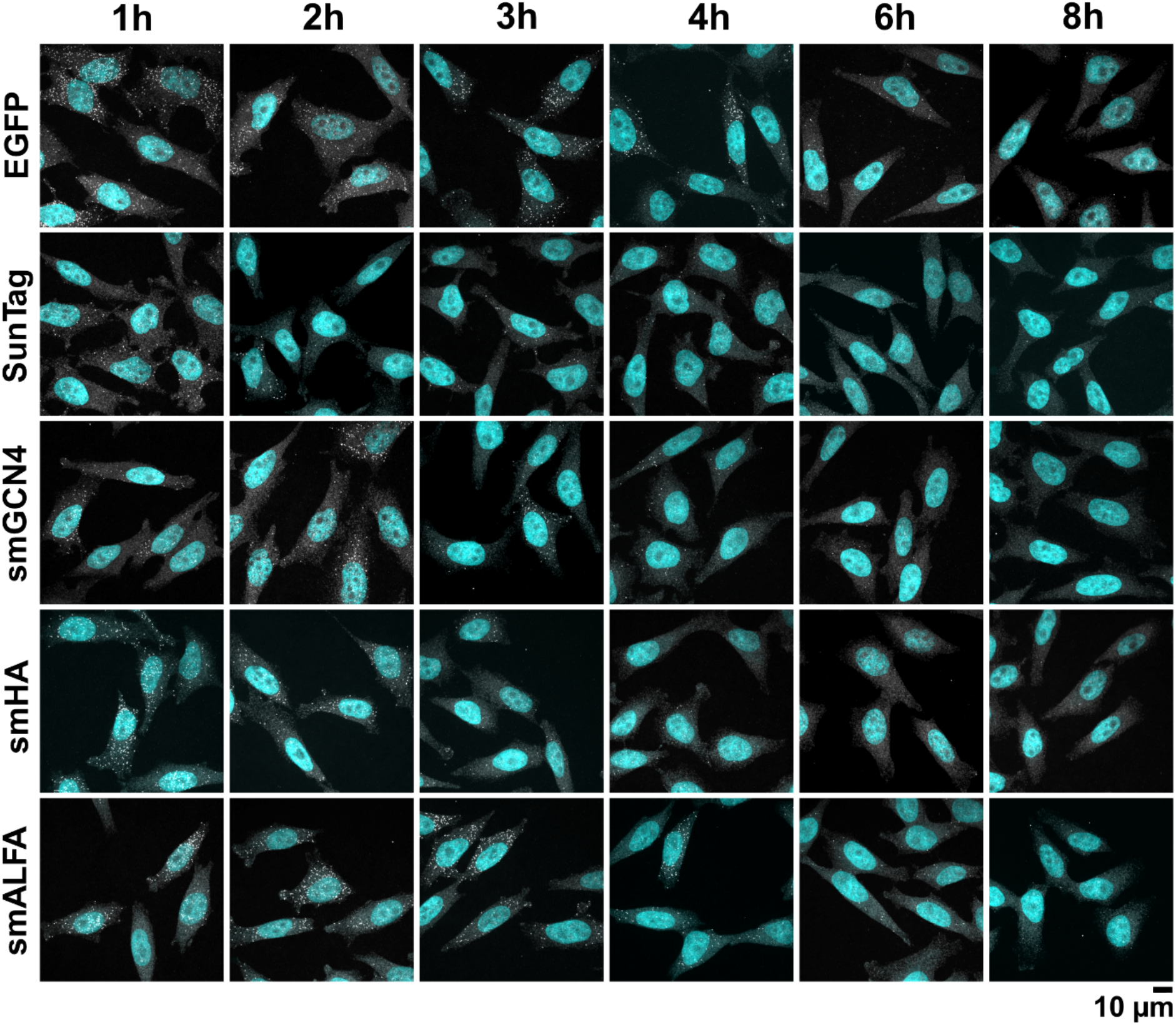
Representative images used for quantification of the number of mRNAs hours after doxycycline induction was removed to assess mRNA stability. A single plane from the center of the cell is shown. Cyan – DAPI, grey – mRNA.

**Supplementary Figure 4.**
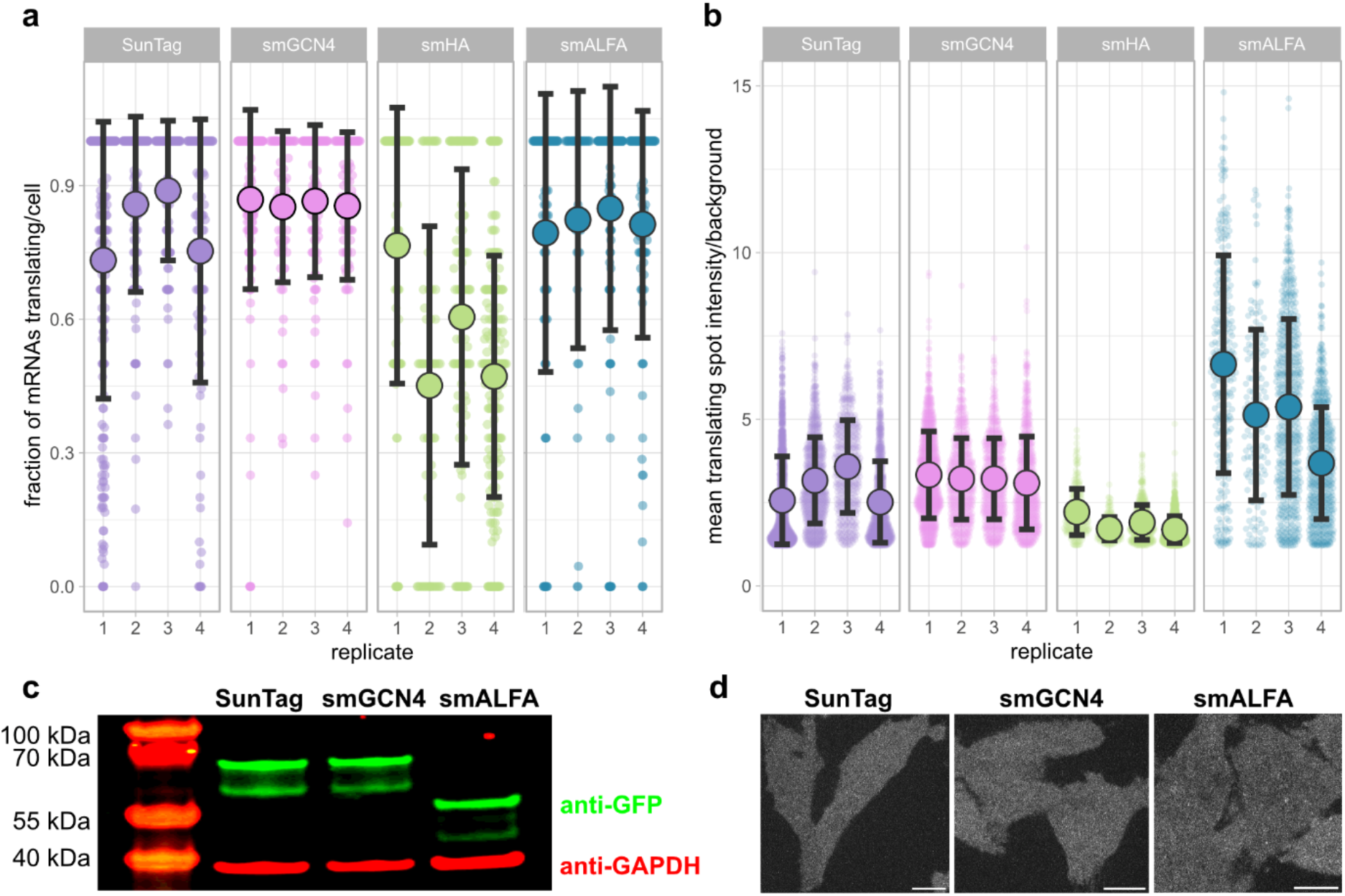
Live-cell single-molecule imaging and protein expression analysis. **a**, Individual replicates showing the fraction of translating mRNAs. **b**, Individual replicates showing the mean intensity of translation spots. **c**, Western blot comparing expression levels of scFv-GFP and ALFA nanobody in the three reporter cell lines. GAPDH serves as a loading control. **d**, Representative images on uninduced cells with the background nanobody-GFP fluorescence. Scale bar: 10 μm

## Notes

### Competing Interest Statement

The authors have declared no competing interest.

